# A definition of song from human music universals observed in primate calls

**DOI:** 10.1101/649459

**Authors:** David Schruth, Christopher N. Templeton, Darryl J. Holman

## Abstract

Musical behavior is likely as old as our species with song originating as early as 60 million years ago in the primate order. Early singing likely evolved into the music of modern humans via multiple selective events, but efforts to disentangle these influences have been stifled by challenges to precisely define this behavior in a broadly applicable way. Detailed here is a method to quantify the elaborateness of acoustic displays using published spectrograms (*n*=832 calls) culled from the literature on primate vocalizations. Each spectrogram was scored by five trained analysts via visual assessments along six musically relevant acoustic parameters: *tone, interval, transposition, repetition, rhythm*, and *syllabic variation*. Principal Components Analysis (PCA) was used to reduce this multivariate assessment into a simplified measure of musical elaborateness. The resulting “acoustic reappearance diversity” index simultaneously captures syllabic variation and spectral/temporal redundancy in a single continuous variable. The potential utility of this index is demonstrated by applying it to several social and habitat-based theories of acoustic display origins. Our results confirm that primate species living in small, monogamous groups have song-like calls, while forest habitat had a less pronounced association.

## Introduction

Musical behavior, or elaborate acoustic display (e.g. song), has independently evolved in several vertebrate [1] and some arthropod [2] clades. However, the historical selection pressures that gave rise to this behavior, and its current evolutionary function, are less well determined. Delineating the emergence of human music, for example, is challenged by its acoustic ephemerality and a paucity of artifacts—although fossil musical instruments have been unearthed [3]. Consequently, we have few clues available to resolve if human music is truly novel or merely an evolutionary continuation of the song-like calls of non-human primates such as those of the lesser apes. Alternatively, researchers can leverage statistical tools to investigate ultimate evolutionary function and mechanism by using behavioral data among living organisms [4].

A number of explanations have been proposed for the evolution of musicality. The primary habitat-based theory, the acoustic adaptation hypothesis (AAH) [5–7], makes two predictions about how the structure of flora drives the evolution of the vocalizations of the inhabiting fauna. First, it predicts that low frequency vocalizations will increase as vegetation density increases [5]. This has been previously demonstrated in primates [6]. The second prediction is that there will be more inter-element intervals as vegetation structure becomes more complex [5]. AAH has been modestly supported over the years but its explanation of song is only weakly supported [8].

Theories on a social function of musical behavior lie on a continuum ranging from the affiliative and parental (e.g. lullabies) to the more aggressive and territorial (e.g. loud calls) with mating calls, or “love songs”, lying somewhere in-between [9]. These include: emotion regulation and language acquisition in infancy [10,11], pair bonding [12], sexual advertisement [13,14], group cohesion [15], group selection [16], and coalition signaling [17]. We suggest that these approaches to musical behavior have fallen short in disentangling the problem of function because they lack a quantitative, acoustic features-based definition.

Historically, the critical testing of these theories has involved using extremely broad, binary aesthetic assessments—in the case of social selection [17], or focusing too narrowly on specific features (e.g. call frequency) in the case of habitat selection [6,8]. Few studies have attempted to combine multiple acoustic features of display *signal* to more objectively encompass an essential musicality—efforts at a definition have suffered from ambiguity [18], circularity [4], and imprecision [19]. Although there have been efforts to investigate the relationship between specific acoustic features of music and specific selective forces [14], few have focused on features of musical *performance* [19] and many approaches only study western music *listeners* [14,17,20] a culture where they vastly outnumber performers [21].

We endeavor to construct a neutral formulation of these acoustic features (the signal itself) based mostly on human music universals [22]. We do so by first distinguishing “utterance level” features—those present in every piece or performance—from “conserved” or “common” features— those present at some level in a musical system or culture [23]. Non-vocal modes of generation (e.g. via instruments) and cultural musical contexts (e.g. dance and rituals) are common human universals [24], but they are rare in other vertebrates. These contexts are also not inherently acoustic themselves and might best be reconsidered as co-evolutionary *influences* on acoustic displays. Accordingly, we focus initial index construction efforts on only *structural* universals (e.g. pitch, rhythm, melody, and form) from human music [24]. Contexts can then instead be tested later as potential influences on this independently constructed acoustic based index. Our approach differs from previous work [1,25], in that features need not be uniquely human, just universally so. And we have omitted certain universals studies [26], in order to focus our analysis on the vocalizations produced by the senders (e.g. tone) rather than the audio perceived by the receivers (e.g. pitch) of musical signals.

Evolutionary bioacoustics outside of our own species (e.g. in birds) has historically focused less on rhythm and pitch and has instead focused on more spectrally complex aspects of display. Examples of these approaches include between-song structural consistency measures such as typicality, the similarity of a song’s features with the songs of others, and stereotypy, consistency in one’s own songs [27]. Other, within-song focused signaling features (e.g. song rate, length, and size), relate more to mate choice preferences for output, performance, or complexity [27]. Here we aim to explore aesthetic feature combinations that span multiple, of these broad (within-song) signaling categories simultaneously. Song analysis has historically entailed visual quantification of various putative aesthetic features of possible signaling importance present in spectrograms—plots of spectral energy over time (Fig. 1). Various within-song measures such as: unit consistency [28,29], trill rate [30], repertoire size [31,32], song bout length [33], and complexity [32,34], overlap well with the utterance level universals we use. But while we’re partial to the category of “complexity,” we aspire to transcend its ambiguous connotation by developing a sophisticated and concrete multi-feature index.

**Figure 1.**
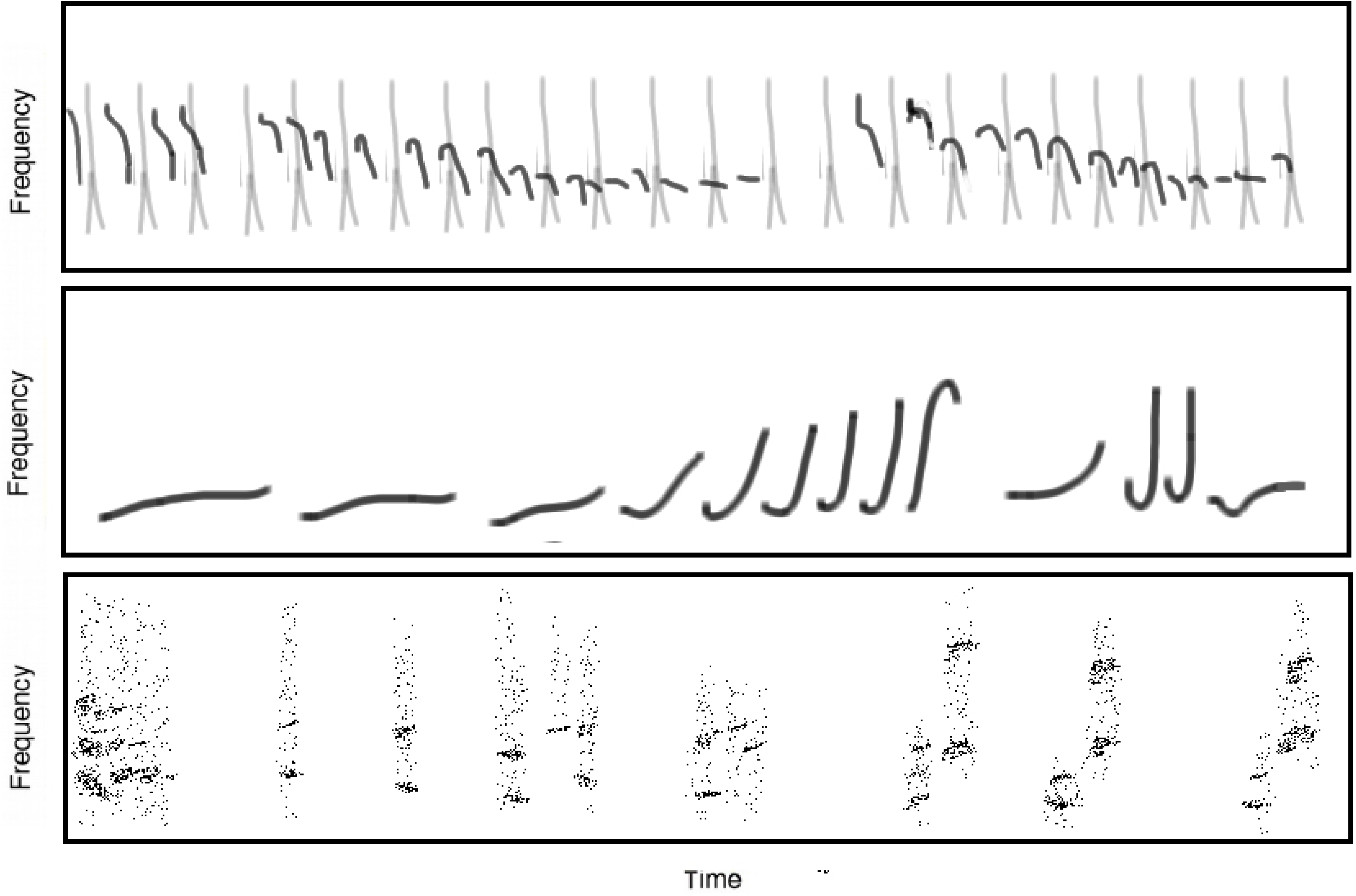
Spectrograms of various elaborate primate calls. Redrawn spectrographic representations of three species calls with corresponding acoustic reappearance diversity scores (averaged across 5 independent visual assessments) formulated as syllables * (Pr(repetition) + Pr(transposition)) where Pr is a proportion**. (top)** *Tarsius spectrum* 4.8 * (0.86 + 0.4) = 6.1; **(middle)** *Nomascus concolor* 5.4* (0.48 + 0.18) = 3.6 **(bottom);** *Lepilemur edwardsi* 2.8 * (0.72+ 0.1) = 2.3. Figures redrawn from Nietsch 2003, Geissmann 2000, and Gosset 2003 respectively (consult the vocalization references listed in the supplemental information file).

The terminology differs between the human and avian bodies of literature, but many of the aesthetic features from both seem to nicely group into two broader categories. First, a similarity among units, and secondly, a diversity among units—measured, for example, via consistency of repeated units or number of different units respectively. These more melodic and form related elements might best be included at the more universal *utterance* level of musical acoustics. The less common system-level universals such as those relating to rhythm and tone (e.g. pitch), may not be as efficient at explaining more diverse dimensions of proto-musicality. Human musical utterances consist of multiple, discrete units (e.g. notes, chords, phrases) that both vary (in pitch, tempo, or texture) and repeat [23,24] (Table 1: utterance). There is disagreement as to whether pitch, a constituent of tone [35], and rhythm are required features at this first, most basic, level of musical organization (Table 1: system). Whereas rhythm and pitch are prevalent in both human music and animal song, they may not be universally common features of all elaborate vocal utterances.

**Table 1.**
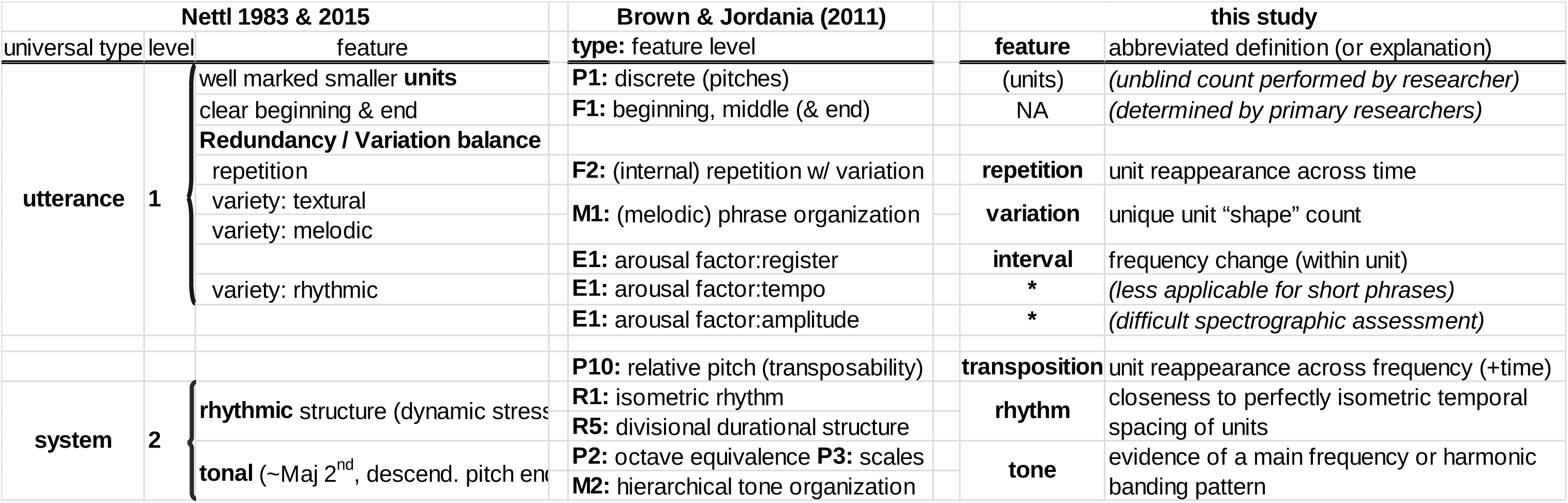
Gradient of music universals across different studies.

There are two main levels of human music universality: utterance and system. The current study attempts to understand utterance level universals. Two previous studies, however, were somewhat ambiguous in determining the level at which the certain features might lie. After the discrete units requirement, repetition and variation are most unarguably universal at the utterance level, and pitch and rhythm are included at the system level, with transposition lying somewhere in-between. ‡Note that while we group Brown and Jordania’s “discrete pitches” with utterance level universals, it’s primarily due to the match-up with Nettl’s “discrete” (not because of “pitch”). While we recognize “discrete pitches” as being ranked highly by Brown & Jordania, we argue for “discrete” as fitting best at the utterance level and “pitched” as fitting best at the system level. *Note additionally that variation in duration, tempo, and amplitude were not tabulated for this study.

To resolve this debate, we developed an impartial formulation by collecting spectrograms of vocalizations from 55 primate species and then scored them on six musically relevant acoustic parameters—at both the *utterance* and *system* levels. We then performed a principal components analysis (PCA) informed variable reduction on these six acoustic feature scores. The contrasting utterance-level acoustic music universals of syllabic diversity and reappearance were retained and combined into a univariate measure of proto-musicality that detects song-like elaborateness from any acoustic utterance. The resultant “acoustic reappearance diversity” index is defined as the expected number of unique spectral shapes or “syllables” that reappear within a call (either by repetition or transposition). We demonstrate the utility of this metric by applying it to key ideas from the two theoretical bodies mentioned above: both adaptation to habitat acoustics and selection based on social influences.

## Data collection and analytical methods

### Vocalization data collection

As an alternative to analyzing raw audio recordings, which are often unavailable, we used published spectrograms: plots of acoustic energy where *x*=time and *y*=frequency (Fig. 1). We sampled spectrographic studies from nearly all families in the primate family tree, where each vocalization collection was individually culled and classified by primatologists focusing on select species. We primarily focused on collecting *continuous* data, from spectrographic vocalization repertoires (for 62 species), and only secondarily on *categorical* call type data (e.g. *loud call, long call, chorus, song, duet*) from text descriptions of vocalizations (for 199 species) [6,36]. The spectrographic studies focused on individual species and were all published in English before 2014. The categorical data (e.g. name, type, and context) were additionally used to verify the multivariate analysis on the variables derived from the spectrographic dataset.

We searched for publications meeting the above criteria by querying on-line search engines (ISI Web of Knowledge and Google Scholar) to locate these vocal repertoires for the quantitative scoring analysis. Initially this involved hand-entering “vocal* AND repertoire* AND [primate genus]” as an all-field query into the journal article search feature of Web of Science citation index online. Use of this text-based meta-database (limited to title, abstract, and keyword fields), however, could not easily detect presence of “spectrograms” as this particular keyword usually only appears in captions or methods sections. High sensitivity search focus within each genus was discontinued after a sufficient number of species from each were obtained. Additional search efforts were instead redirected toward more sparsely studied corners of the primate family tree using reference cross-checks and review article citations.

In general, studies were catalogs of individual species behavior rather than developmental, experimental, or species comparative studies. Acceptable articles had to include, for each species studied, spectrographic depictions for multiple calls, in order to obtain a variance, estimate of each species’ song index. A primary objective was to obtain “complete repertoire” studies and, as a result, over 2/3rds of accepted studies had more than 10 different calls (*n*=45 species). Some exceptions were made for species with (an) obvious, stand-out display call(s) (e.g. gibbon songs) that were otherwise relatively non-vocal (*n*=5). Some other exceptional non-repertoire focused studies (e.g. long calls, loud calls) were also included (*n*=5). Because the main goal was to let acoustic features predict song-like calls independent of researcher call designation, we did not include any other studies on just a single call type (e.g. contact, food, alarm). A single study (Harcourt 1993) that was neither a full-repertoire nor a loud-call study on the “close calls” of the gorilla was used as studies with a larger variety of calls were not found.

We scanned 61 books and downloaded 67 PDFs to obtain spectrographic vocalizations from more than 80 species and over 300 total leads on possibly relevant studies. Only a single spectrographic study for each species was used in the data-set, so that some studies of the same species were removed (*n*=53). In these cases, we retained the higher quality studies: those with more vocalizations described, more modern recording and analysis tools, higher quality spectrograms, more sophisticated call classification technique, or that were more recently published. The final collection of spectrograms was extracted from 58 sources resulting in 1297 different spectrograms for 61 species representing 40 genera.

For 44 studies in electronic format, images were obtained as screen captures at 100% zoom. For the remaining species, we scanned spectrograms from printed articles at 300dpi as grayscale 8-bit depth bitmaps to provide similar resolution. We also used image editing software to manually clean and standardize the spectrograms by removing axes, labels, and any annotative markings.

Vocalizations were grouped into 842 species-specific note, phrase, and song types as assigned by the original authors. We included as separate vocal types both single unit and repeated unit vocalizations, if the primary authors had also done so. Ten vocalizations (from three different studies) did not meet the minimal study acceptance criteria above, leaving 832 scored vocalizations (corresponding to 1287 spectrograms from 55 sources).

### Spectrogram scoring

We used simple human music universals [23,24] and the principles of acoustics [37] to guide us in selecting a total of six structural features as scoring parameters: variation, repetition, transposition, rhythm, tonality, and interval. Spectrographic interpretations of definitions [35,38,39] used (see Table 2) are abbreviated as follows:

**Table 2.** Vocalization spectrogram component definitions and scoring key.

| **acoustic feature** | **definition** | **spectrographic** | **scoring** |
| --- | --- | --- | --- |
| tone | a steady, regular, or periodic sound characterized by its pitch (or perceived frequency), intensity, duration, and timbre | presence of a clean, high-contrast fundamental frequency and/or harmonics | low={noisy, pixelated, grey}<br>high={clean, clear, black & white} |
| interval | the difference between two (successively sounding) tones. The ratio between two sonic frequencies | the presence of sloping or curving (rather than static) fundamental frequency within units | low={flat or noisy}<br>high={sloped, jagged or curvy} |
| rhythm | a regular recurrence or pattern in time--a movement marked by the regulated succession of strong and weak elements, or of opposite or different conditions | the regularity of spacing between units over time | low={unpredictable horizontal spacing of units}<br>high={evenly spaced repeats, scale or syllables} |
| repetition | restatement [over time], such as the restatement of an utterance, phrase, or theme. | reappearance of a syllable at the *same* frequency across time | low={isolate, unique}<br>high={all units match many other units horizontally} |
| transposition | moving a (collection of) note(s) up or down in pitch by a constant interval | reappearance of a syllable at *different* frequencies over time | low={flat progression}<br>high={all units match other units after shifting vertically and horizontally} |
| syllable(s) | a unit of organization for a sequence of speech sounds. Typically made up of a syllable nucleus (most often a vowel) with optional initial and final margins (typically, consonants). | a count of the number of distinct unit shapes within a call | low={one unit shape type}<br>high={many unit shape types} |

1. **tone:** the presence of clean harmonics with distinct, horizontally-parallel bands
2. **interval:** a sloping, jagged, or curving, rather than static, fundamental frequency
3. **rhythm:** a regular recurrence or pattern of [vocal] units over time
4. **repetition:** similarity in [vocal] units repeated across time
5. **transposition:** similarity in [vocal] units of different frequencies (and at different times)
6. **variation:** number of distinct [vocal] unit types or shapes (“syllables”) within a call

Observers were trained for one hour on feature definitions and how they can be identified and quantified from spectrograms. Manual scoring was performed blindly without reference to the species. Vocalizations were scored for each of the six musical features in a random order of species. Each of the six features was scored on a scale of 0 (lowest) to 10 (highest), except for variation which was scored as a count of unique syllable shapes. Vocalizations with multiple spectrograms (*n*=221 or 26.6%) were separately analyzed and then averaged to calculate the final measure for that vocalization type. This matrix of ordinal scores was then averaged across the individual scorers. Finally, for the PCA analysis, these scores were scaled to continuous values between 0 and 1.

Listed here are our six acoustic music universals, their definitions [35,38,39], and a spectrographically relevant interpretation for scoring purposes. The first five dimensions were scored on a scale of 1 (lowest) to 10 (highest), while syllable was scored as a count of different spectral shapes.

### Principal components and dimension reduction analysis

We used PCA as a guide in reducing the acoustic feature scores from six to just three variables that could then be combined into a single multivariate ‘elaborateness’ index. In this data set, for example, repetition and rhythm are highly correlated with each other as are tone and interval. These two variable pairs are therefore strong candidates for reduction where one variable from each pair is kept as a proxy for both variables in the pair. The end goal of this reduction is to both eliminate redundancy and gain access to statistical analysis programs and functions that require a univariate parameter as input. Using PCA to inform a dimensionality reduction also has several additional advantages ranging from alleviating visualization issues to addressing multicoliniarity of variables [40].

PCA is an exploratory statistical procedure that orthogonally transforms a dataset (of *n* observations on *p* possibly correlated variables) into a set of linearly uncorrelated principal components [40]. In this case, *p* corresponds to six music universal feature scores and *n* equals 829 primate vocalizations. The loadings (i.e. weights, or correlations) of the original *p*=6 variables with each of the components, are a useful way to systematically translate between the original variables and these main variance-explaining best-fit lines. The loadings were used as a guide in selecting a subset of variables that encapsulate most of the variation. This involved selecting the variables with the highest loading (α_0_), or contribution, in the retained components (α_0_ > 80%) and discarding those variables associated with low eigenvalue (λ_0_ < 0.7) components [41].

### Index development, verification, and demonstration

We use a probability argument to develop an index that most efficiently captures acoustic elaborateness at the utterance level. We also use theoretical arguments—invoking norms from avian bioacoustic research, human music history, and ethnomusicological works [23,24]—to support the acoustic feature selection. For verification we performed Mann-Whitney U tests and Pearson’s Rank of the index against established call names and contexts. We also illustrate the utility of the resulting index by examining theories of song and music evolution.

## Results and discussion

### PCA results

The results of the PCA (Table 3) suggest that PC1 (the best-fitting variance-minimizing line) is one that delineates along a continuum from signal-rich, song-like calls to acoustically noisy and single unit calls (Fig. 2 a and b respectively). We hereafter refer to this as the “signal content” component. All loadings in this component are negative suggesting that all six features contribute to explaining the signal content component and are helpful in assessing acoustic musicality. This first signal content component explains 43% of the variance (Table 3).

**Table 3.** Results of the principal components analysis of music universals on primate calls.

| Acoustic Feature | Component |  |  |  |  |  |
| --- | --- | --- | --- | --- | --- | --- |
|  | 1 | 2 | 3 | 4 | 5 | 6 |
| syllables | -0.40 | -0.03 | -0.19 | **0.87** | -0.17 | 0.08 |
| repetition | -0.46 | **0.53** | 0.07 | -0.23 | 0.04 | 0.67 |
| transposition | -0.26 | -0.42 | **-0.79** | -0.33 | -0.05 | 0.13 |
| rhythm | -0.52 | 0.40 | -0.09 | -0.17 | 0.09 | **-0.72** |
| tone | -0.39 | -0.39 | 0.46 | -0.22 | -0.66 | -0.03 |
| interval | -0.37 | -0.48 | 0.33 | 0.02 | **0.72** | 0.05 |
|  | **1** | **2** | **3** | **4** | **5** | **6** |
| Loadings (eigenvalues) | 2.55 | 1.29 | 0.83 | 0.71 | 0.49 | 0.13 |
| Proportion Variance | 0.43 | 0.22 | 0.14 | 0.12 | 0.08 | 0.02 |
| Cumulative Variance | 0.43 | 0.64 | 0.78 | 0.90 | 0.98 | 1.00 |

**Figure 2.**
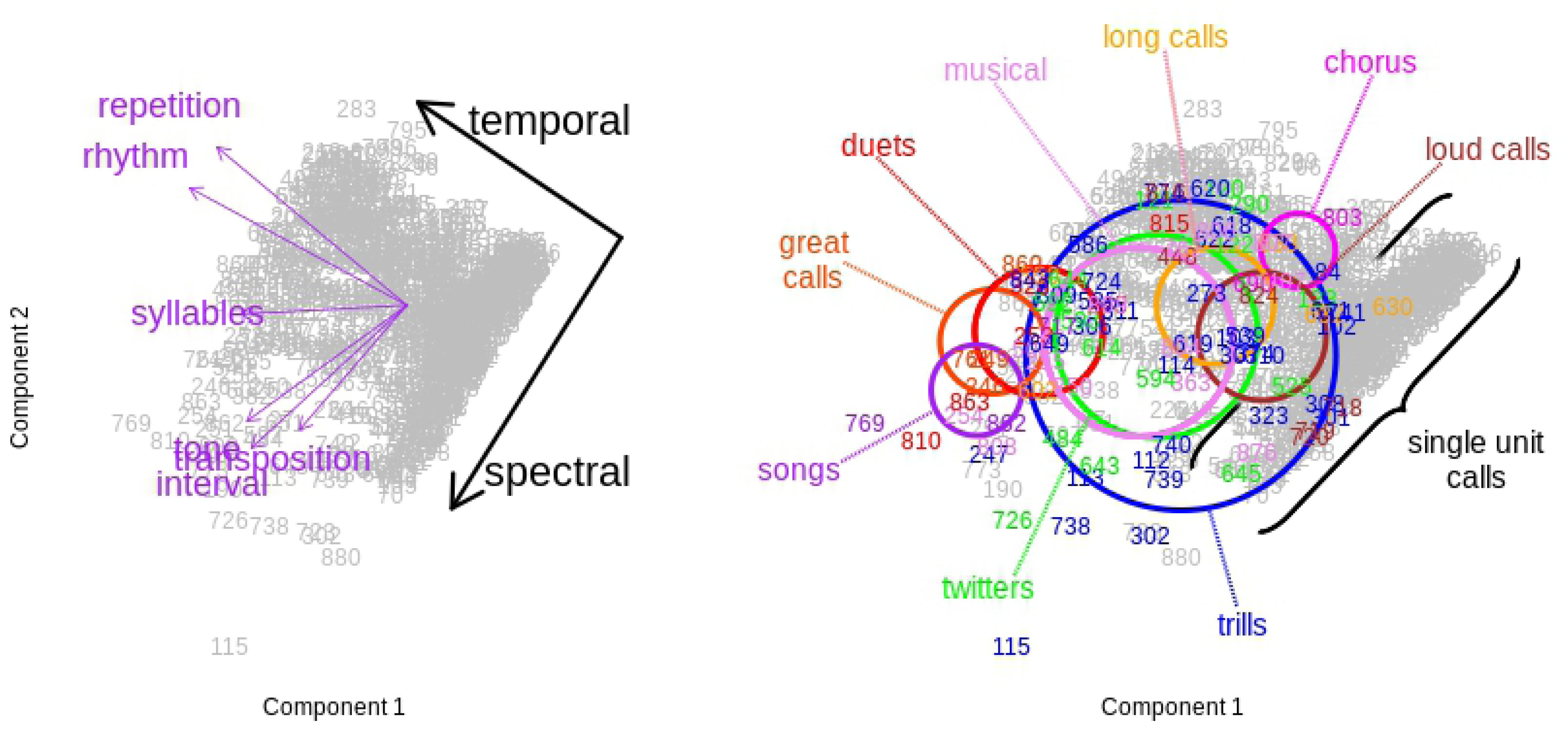
Principal Components Analysis [PCA] of six acoustic display aspects of primate calls. PCA on six acoustic music universals (tone, interval, and rhythm, repetition, transposition, and syllable count) where each numbered point above represents one of 823 unique primate vocalizations. (a) Each of the six arrow-head coordinates represents the loadings (contributions) of each of these acoustic feature scores towards PC1 and PC2 (also see Table 3). The three distinct clusters formed by these PC loading coordinates, suggests a possible reduction in dimensionality down to just three proxy measures – a diversity measure: syllable count (left) and two redundancy measures: temporal (top) and spectral (bottom) **(b)** Using the same underlying numbered point positions of calls, primary study author determined call types labels and rings (color online) point out their approximate clustering ranges, scaled to relative number of calls. The plot suggests that song like calls (b: far left) are distinctly more signal rich than the (non-overlapping) long calls, loud calls, or choruses (b: right).

The Principal Components Analysis of human-music acoustic universals *(p*=6) applied to primate calls (*n*=826), suggests that repetition, transposition, and syllable count are the most explanatory of overall variance. The feature score loadings (top table) contains each features’ correlations with the components: PC1 (signal content) explains 43% of overall variance and indicates all six parameters contribute to a call’s signal content; PC2 (degree of temporal versus spectral redundancy: 22% of total var.) highlights the high loading of repetition (53% corr.). PC3 (14% of var.) and PC4 (12% of var.) have top loadings of transposition (79% corr.) and syllable count (87% corr.), respectively, of feature score correlation with each component. Loadings which were highest in absolute value both across features and across components were highlighted in bold (PC1 had no such value).

The second component, that minimizes the variance between the first component and the residuals of that component’s fit, is one that differentiates between types of redundancy: temporal versus spectral (Fig. 2 a and b: top and bottom respectively). The highly correlated time domain measures of rhythm and repetition both have positive loadings and the spectral domain measures of tone, interval, and transposition all have negative loadings along PC2.

A pronounced inflection point in eigenvalues between these first two components (PC1: signal content: λ=2.56 and PC2: redundancy: λ=1.3) and the rest suggests that we might focus primarily on the former and less on the latter. The third and fourth components, however, do explain a good proportion of the overall variance—raising it 25.5% from 64% to 90%—and the eigenvalues (λ_0_) are all above 0.7 and suggest retention [41]. These two components are harder to interpret than the first two (signal content and redundancy type), but the loadings correlations, of each parameter with each component, are informative. The single highest loadings for each of these two components are, interestingly, transposition (79% loading correlation) and syllable (87% loading correlation). They explain 13.8% (PC3) and 11.7% (PC4) respectively of overall variance—after 42.5% (PC1) and 21.6% (PC2).

Syllable count is the most unambiguously neutral in PC2 (redundancy) and clearly collimate with PC1 (signal content) suggesting it could be an efficient indicator of complex calls. As mentioned above, it was also the highest loading feature in the 4th component—one which explains 12% of the variance of the overall dataset. Syllable diversity’s prominence is not that surprising as its analog (repertoire size) is a commonly used metric for display quality in avian acoustic research [42,43].

Repetition and rhythm had similar loadings in PC1 and PC2 (Fig. 2a) suggesting a collapsing of them into a single variable to reduce collinearity. Rhythm was indicated as being important, but it was excluded from the index due to its high association (72%) with discarded PC6 (λ=0.13). Only one of these was kept as either of these two alone could serve as a rough proxy for time-domain redundancy. Repetition is more elemental (as it is often a prerequisite for rhythm) and is thus further justified for retention in the index. We offer additional rationale below in arguing for rhythm’s proper classification as a musical *system level* universal (also see Table 1).

The PC1 and PC2 loadings for tone, interval, and transposition similarly overlap with each other in the PCA analysis (Fig. 2 a and b bottom) and could be reduced to a single representative non-co-linear variable representing frequency domain redundancy. Interval was difficult to properly assess as were other (unmeasured) emotive/arousal universals (e.g. tempo and amplitude variation). As it had the highest loading with the discarded fifth component (λ=0.49), interval was ruled out. Pitch, like rhythm, has an unclear position in the gradient of musical universality somewhere between utterance and system level universals [23,24], and it’s possible that tonal (pitched) units should not be categorically required in an utterance level definition (Table 1). Transposition, with its high loading on the third component, was selected to serve a proxy for both pitch and interval.

### Towards a univariate quantitative index

The topic of musical quality can be quite polarizing [4,21] and features such as pitch (e.g. tonal versus atonal music) and rhythm (e.g. melodic versus rhythmic music) are likely candidates for inciting disagreement. Perennial controversy surrounding these features hint that they don’t necessarily both belong in a universal definition. Our PCA results correspondingly suggest we can be less concerned with these two features as they can be proxied by the spectral and temporal redundancy measures of transposition and repetition. These features’ components together explain over a third of the total variance (where as the rhythm associated component explains less than three percent).

Acoustic display has more simply and broadly been described as an emergent balancing of ‘[ritualization with innovation]’, or “an unusual combination of order and chaos” [14], of “redundancy balanced by variety” [23], or “internal repetition with variation” [24]. Variety unquestioningly provides the combinatoric uniqueness underlying musical novelty and interest. But its counterpart, repetition, though it provides baseline temporal acoustic scaffolding, still remains a relatively unsolved mystery in musicological research.

We need only include this minimum set of acoustic universals, as we are most interested in detecting song at the most abstract levels. And we could require the two simplest and yet nicely contrasting and balancing features of redundancy (proxied by *consistency*) and variation (proxied by *size* or *complexity*) of syllables within an utterance—especially given the (avian/human) quality metric overlap discussed in the introduction above. The PCA nicely corroborates this theoretical argument for a simple inclusion of just these few non-collinear variables. However, we still need to further quantitatively combine them, if we are to obtain a single outcome measure of acoustic elaborateness. Below, we provide the mathematical rationale for adding the two reappearance probabilities together and then multiplying the result by syllable count.

These two (within-utterance) features can be quantitatively defined as follows: *variation as* a count of the number of distinct syllables and *redundancy* as reappearance of syllables across time— either at the same frequency, in the case of repetition, or at different frequencies, in the case of transposition. Mathematically, we need to determine which operations to use when combining these together. As for combining repetition and transposition, we can re-purpose the addition rule of probability theory [44] that states that for two events, A and B:

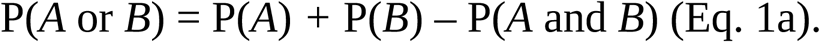

The last term can be set to zero due to mutually exclusivity [44] of the repetition and transposition of any given vocal unit. That is, it’s impossible to both repeat, in time, and transpose, in frequency, a unit across an entire call. And since these two feature scores also happen to be easy to scale into probabilities, as they are already recorded on a scale of 1 to 10, the probability of unit reappearance as the sum of the two terms can be written like so:

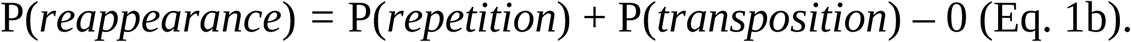

For integrating this new reappearance probability into our index, we can model the index (which requires both unit reappearance and syllabic diversity) as an expectation [44] written like so:

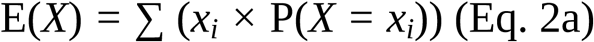

where X is a random variable that serves as an indicator of reappearance. It is a binary (yes or no) variable that answers the question: does this unique syllable [i] occur elsewhere in the utterance? The probability term can be removed from the summation because it is uniform across the entire call (scoring was assessed on entire calls and not individual units). The equation, within the context of this study, then simply becomes the count of unique syllables times the overall probability of syllable recurrence within the utterance:

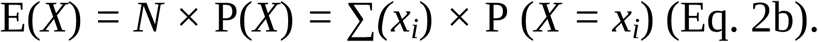

Rewritten with the full names of the two main components, this expectation looks like:

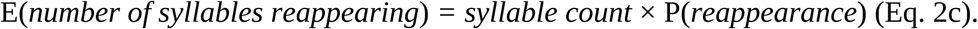

This use of multiplication is an elegant and mathematically certain way to require that each of these elements co-exist within every musical utterance; *multiplication* of the two individual feature scores of syllable and reappearance guarantees a score of zero if either feature is scored as zero (Equation 2c).

### Corroboration of the index

We demonstrate use of the acoustic reappearance diversity index by examining its correlation with vocalization categories and contexts. The appropriateness of the composite index was suggested by its assignment of relatively higher values to vocalizations categorized as *display* (Mann-Whitney U test, *n*=829, *W*=3581, *p*<0.0001) or those described as *song, duet, trio, chorus, great, music, scale, coda, intro*, or *interlude* (Pearson’s rank, *n*=829, *r*=0.49). Visual evidence of the latter of these correlations is available by inspecting the overlay of these song names on the PCA plot (Fig. 2b). The correlation between higher acoustic reappearance diversity index values with classifications such as ‘duet’ or ‘song’ (Wilcox-test: *n*=58, *W*=91, p<0.007; Fig. 3) verified this composition of features in the composite score. Similarly, higher scores also associate with primary author determined call contexts such as “display” and “sociosexual” (Fig. 4). These scores are univariate, continuous, blindly scored, and conform to expert-determined names and contexts.

**Figure 3.**
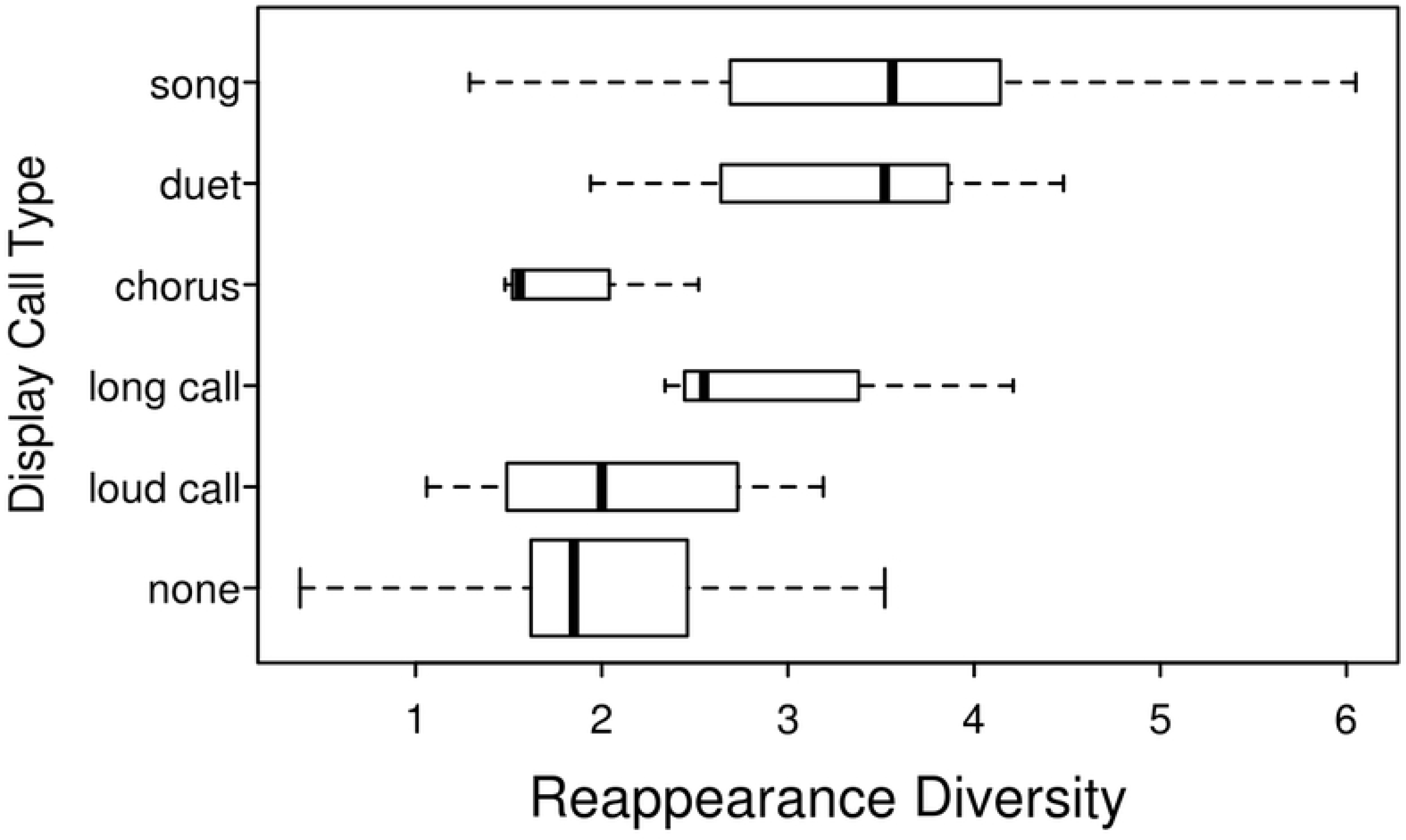
Acoustic reappearance diversity scores versus display call type. Acoustic reappearance diversity is a univariate measure of the expected number of times a unique syllable reappears within a single call or song. This measure has a high correspondence with duet or song calls versus other types of calls.

**Figure 4.**
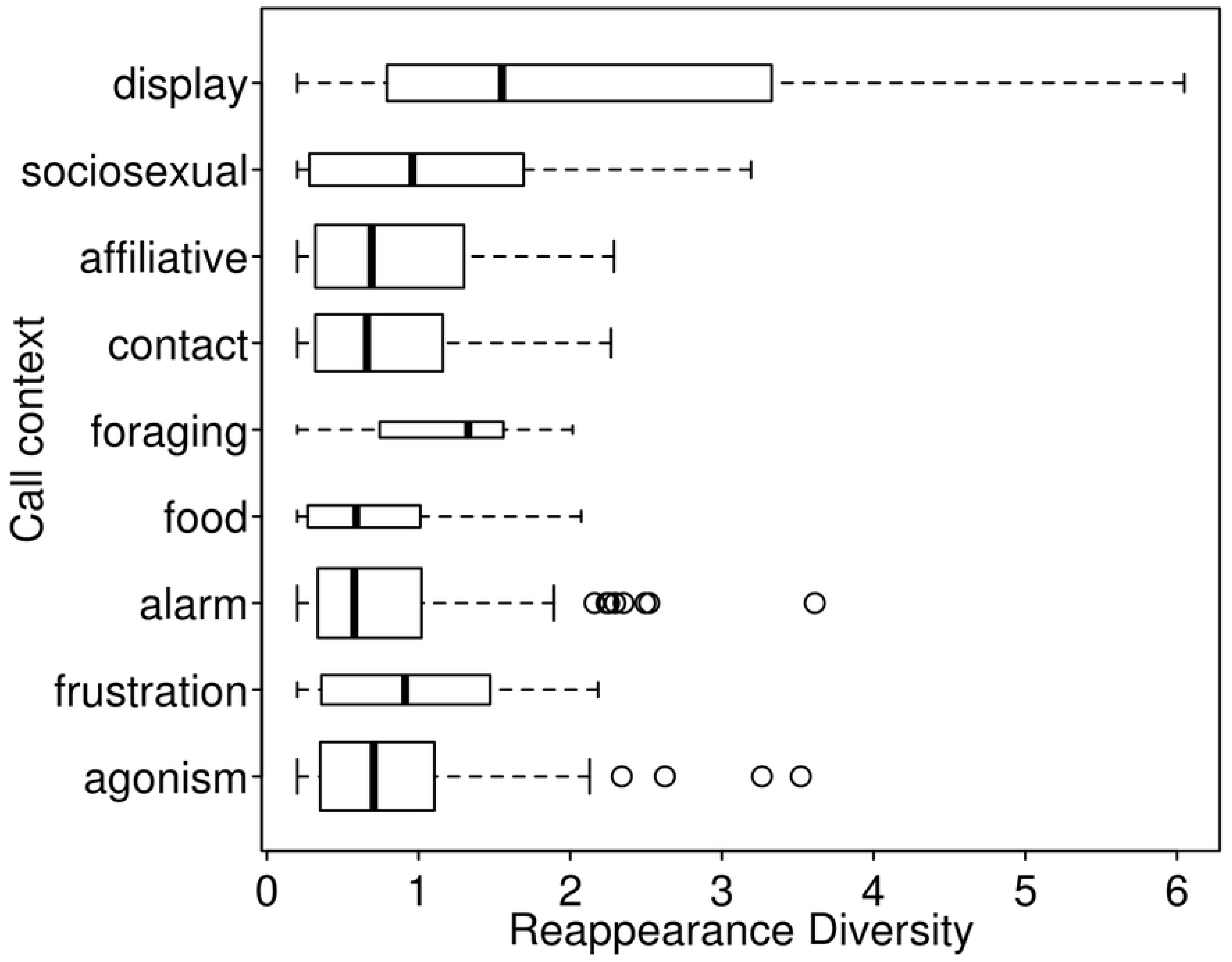
Acoustic reappearance diversity scores versus call level controls (context). Display calls are the primary vocalization context which appear to strongly associate with higher acoustic reappearance diversity. This suggests that the reappearance diversity measure could serve well as an indicator of elaborate display calls.

These results seem to corroborate our index formulation, but there admittedly exists potential western bias in that both the primary researchers (naming and classifying calls) as well the five trained scorers (using English definitions) are mostly culturally western and primarily English speaking. Thus there remains bit of circularity in validating an index built upon western feature definitions, scored by mostly western students, using western researcher determined call names. Although future studies could include more scorer diversity, we don’t anticipate a significant impact on the results as we attempted to be as objective and blind as possible. And although we have admittedly not completely avoided all forms of definitional circularity, we have tried our best to minimize these self-fulfilling influences.

### Testing habitat acoustics and social effects using a species-level index

We used a single index value for each species to explore questions on music and song origins. A box plot of all species in the study illustrates the range of possible scores within a species from which the top score was selected (Fig. 5). The maximum score for each species was used, because we are interested ultimately in the highest level of possible performance in the display calls of species. This “max acoustic reappearance diversity” index formulation showed negligible correlation with many possible species or study and species level predictor variables, but significant exceptions such as habitat, monogamy (Fig. 6) and group size (Fig. 7) are discussed hereafter.

**Figure 5.**
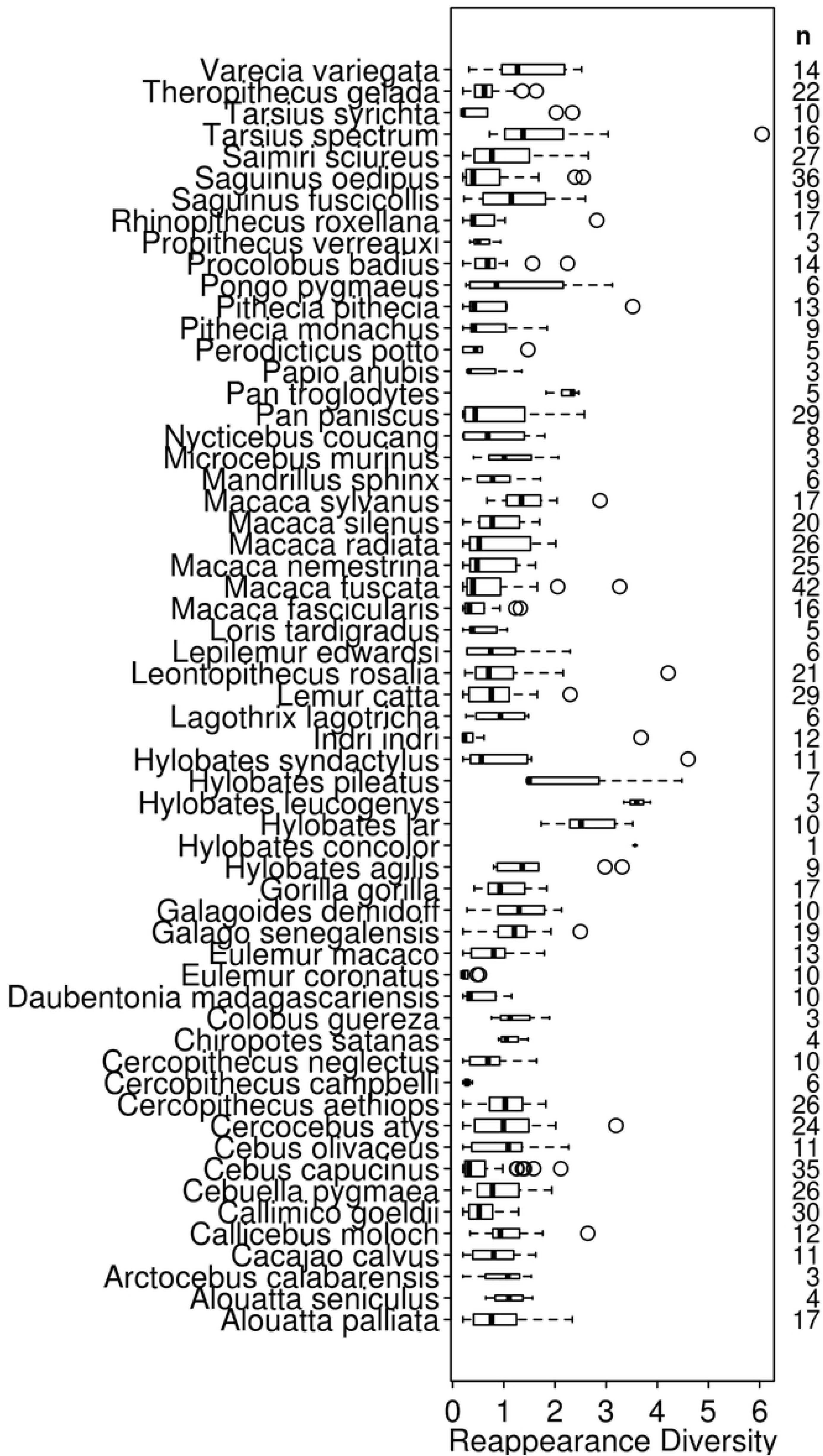
Species level reappearance diversity scores distributions. These box plots demonstrate how species can have a great array of calls (total count of calls in repertoire listed under “**n**” on right hand side) whose reappearance diversity scores are quite low but have a stand-out call (e.g. *Indri’*s song) which scores exceptionally high.

**Figure 6.**
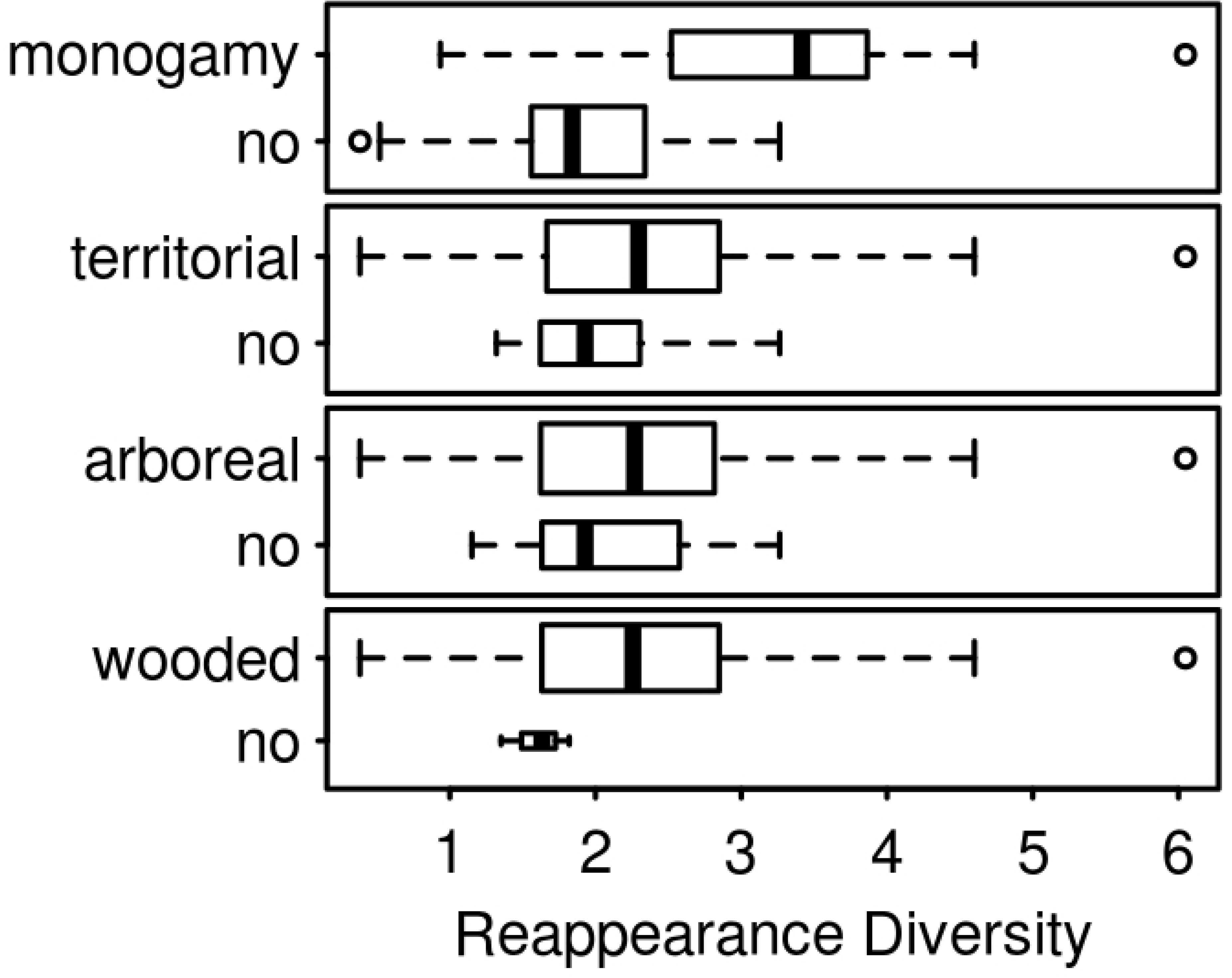
Acoustic reappearance diversity scores versus socioecological controls. The emergence of musical behavior has many explanations ranging from territorial defense to an acoustic adaptation to pair-bonding. While the score disparity between wooded and non-wooded habitats lends support to the acoustic adaptation hypothesis only monogamy appears to have a strong association with acoustic reappearance diversity.

**Figure 7.**
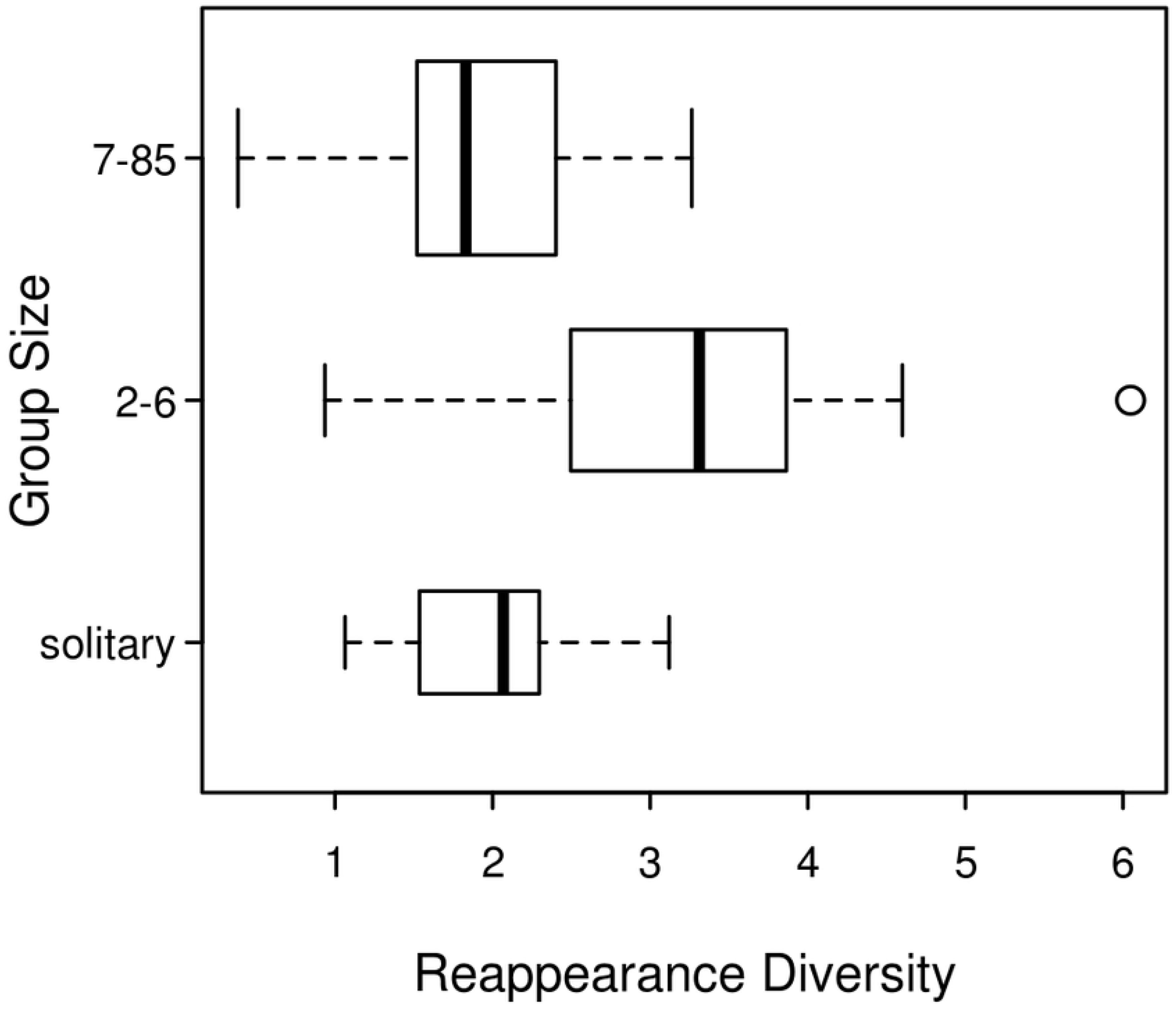
Box plots of reappearance diversity scores versus typical group size per species. Plotting group size categorically, higher acoustic reappearance diversity scores predominate in small group size species (e.g. monogamous, duetting primates such as gibbons, tarsiers, and Callitrichids)

The first theory we considered was the acoustic adaptation hypothesis. We were not able to test the fundamental-frequency based component of AAH as we had focused on tabulating more relativistic song-like parameters. The data presented here do, however, suggest mild support for the second part of the AAH regarding inter-element intervals. Species living in wooded habitats had a call with 0.75 (on average) more reappearing syllables (*t*=3.77, *df*=9.74, *p*=0.004) which seems to suggest that changes in habitat acoustics could moderately effect song elaborateness (Fig. 6). Richer variables types, however, beyond our merely binary arboreality measure, are needed to explore effects of higher dimensional wooded habitats and associated behaviors.

The second theoretical body we tested concerned effects of sociality on elaborate acoustic display behavior. Our index and data-set support the prevailing view of monogamy as being an important coevolutionary factor (Fig. 6). Monogamous species had 1.2 (on average) more reappearing syllables for their most-elaborate call. We found less support for a strictly positive linear correlation with group size, but our metric does indicate that species living in small-sized groups (*n*=2 to 6) possessed more song like calls (Fig. 7). Compared with large groups or solitary species, small groups had almost 50% more reappearing syllables (on average) in their most-elaborate call (*t*=3.58, *df=*20.1, *p*=0.002). Although testing all social influences is out of the scope of the present work, our approach here suggests that smaller, more intimate groups merit further, more detailed investigation.

## Conclusion

Song appears to be detectable using the combination of a simple (syllabic) diversity measure and a redundancy proxy measure—one that captures either spectral or temporal patterning. Our PCA determined formulation corroborates a human music universals scheme [23] that emphasizes repetition and variation of discrete units as foundational. Correspondingly, the definition of animal song might not need to be complicated by pitch and rhythm as feature requirements—although “acoustic reappearance diversity” could be re-construed and rarefied to capture them anyway (see Fig. 8). Furthermore, we have similarly argued that music should not be delimited by non-acoustic universals such as mode of generation or context, despite the fact that many cultures consider elaborate acoustic display as inseparable from dance for example.

**Figure 8.**
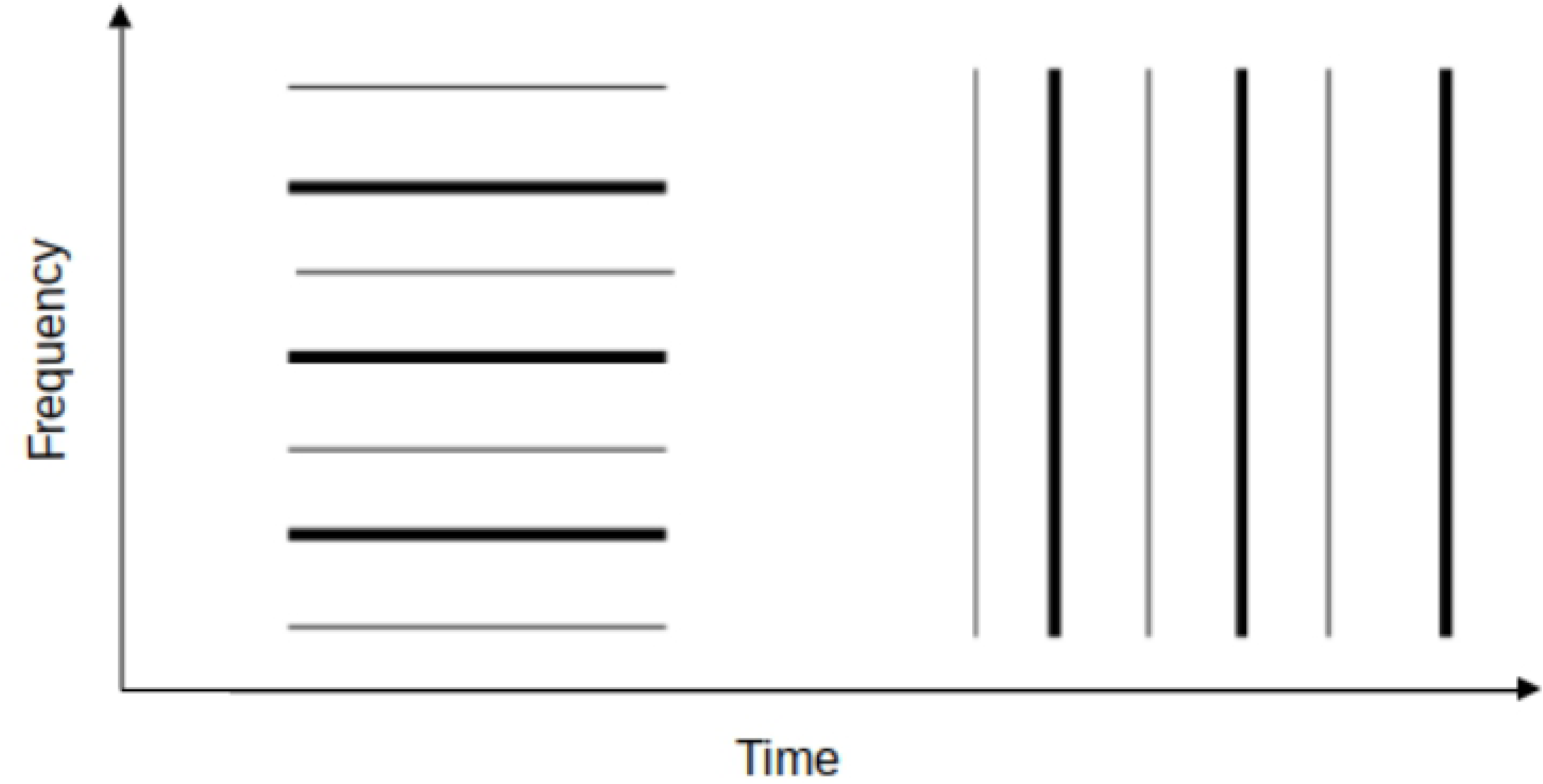
Acoustic reappearance diversity (in amplitude) also captures harmony and rhythm. A highly-simplified illustration of how “acoustic reappearance diversity” could be construed as a general enough construct to encapsulate aspects of both pitched and rhythmic musicality, despite the fact that it was not formulated using either. A pitched matched harmonic sound (left) with two different overlapping harmonic series (the higher frequency tone is bolded as it overlaps with the harmonics of the lower frequency tone one octave below it). A rhythmic pattern (right) with stresses, in bold, every other beat. This illustration is only a very simple demonstration of how our index could be expanded beyond the syllable or utterance level to incorporate higher system level universals. In the example above, it is expanded to include reappearing diversity of ***amplitude*** across both frequency (left) and time (right). The reappearance diversity index could also conceivably be expanded to include much higher-order and complex attributes such as musical motif patterning or song repertoire typicality.

Less simplistic, beyond utterance-level definitions of music (e.g. those including rhythm and harmony), however, should prove to correspond with later co-evolutionary selection pressures such as coordinated group action. These suggested improvements on our work will be useful in assessing influences of ulterior events in the evolution of hominid musicality—those in the last 5 million years. It seems, however, that these large group social contexts were likely not necessary as initial drivers of ancient primate song co-evolution (see chorus opposite of song in Fig. 2). Instead, coordinated musical display of modern humans likely evolved piecemeal from elaborate duet advertisements of small group living and socially monogamous hominoids, which, in-turn, evolved piecemeal from even simpler, more solitary display calls of ancient primates.

## Acknowledgements

We thank Rob Tennyson, Aditya Khanna, Sarah Stansfield, and Jeannie Bailey for scoring spectrograms, Mike Beecher for helpful comments, and Eric Smith for help in data collection and reading of early renditions of this work.

## Author contributions

DMS collected most of the spectrograms, scoring, and control data, with help from DJH, DMS processed and analyzed it, DMS, CNT, & DJH wrote the paper

## References

1. Hauser MD, McDermott J. The evolution of the music faculty: a comparative perspective. Nat Neurosci. 2003;6: 663–668.

2. Pollack GS. Sexual Differences in Song Recognition in Crickets. Am Zool. 1980;20: 853–853.

3. Conard NJ, Malina M, Münzel SC. New flutes document the earliest musical tradition in southwestern Germany. Nature. 2009.

4. Fitch WT. Four principles of bio-musicology. Philos Trans R Soc B Biol Sci. 2015;370: p.91.

5. Hansen P. Vocal learning: its role in adapting sound structures to long-distance propagation, and a hypothesis on its evolution. Anim Behav. 1979;27: 1270–1271.

6. Mitani JC, Stuht J. The evolution of nonhuman primate loud calls: acoustic adaptation for long-distance transmission. Primates. 1998;39: 171–182.

7. Morton ES. Ecological sources of selection on avian sounds. Am Nat. 1975;109: 17–34.

8. Boncoraglio G, Saino N. Habitat structure and the evolution of bird song: a meta-analysis of the evidence for the acoustic adaptation hypothesis. Funct Ecol. 2007;21: 134–142.

9. Lehmann C, Welker L, Schiefenhövel W. Towards an ethology of song: A categorization of musical behaviour. Music Sci. 2009;13: 321–338.

10. Dissanayake E. Antecedents of the temporal arts in early mother-infant interaction. In: Wallin NL, Merker B, Brown S, editors. The Origins of Music. Cambridge, Massachusetts: MIT Press; 2000. pp. 389–410.

11. Trehub SE, Trainor L. Singing to infants: Lullabies and play songs. Advance Infancy Research. 1998. pp. 43–78.

12. Geissmann T. Evolution of Communication in Gibbons. Zurich University. 1993.

13. Darwin C. The Descent of Man and Selection in Relation to Sex. New York: Modern Library; 1871.

14. Miller GF. Evolution of Human Music through Sexual Selection. The Origins of Music. Cambridge, Massachusetts: MIT Press; 2000. pp. 328–360.

15. Roederer JG. The Search for a Survival Value of Music. Music Percept. 1984;1: 350–356.

16. Brown S. Evolutionary Models of Music: From Sexual Selection to Group Selection. In: Tonneau F, Thompson NS, editors. Perspectives in Ethology. Boston, MA: Springer US; 2000. pp. 231–281.

17. Hagen EH, Bryant GA. Music and dance as a coalition signaling system. Hum Nat-Interdiscip Biosoc Perspect. 2003;14: 21–51.

18. Cross I. Music, cognition, culture, and evolution. Biol Found Music. 2001;930: 28–42.

19. Kondik K. A Critical Review of Three Theories for Music’s Origin. M.A., Ohio U. 2010.

20. Pinker S. How the Mind Works. New York: W.W. Norton & Company; 1997.

21. Cross I. When? Musical histories. In: Clayton M, editor. The cultural study of music: A critical introduction. New York: Routledge; 2011. pp. 15–27.

22. Mache F-B. Necessity of and Problems with a Universal Musicology. The Origins of Music. Cambridge, Massachusetts: MIT Press; 2000. pp. 472–479.

23. Nettl B. The Study of Ethnomusicology. Chicago: University of Illinois Press; 1983.

24. Brown S, Jordania J. Universals in the world’s musics. Psychol Music. 2013;41: 229–248.

25. McDermott J, Hauser MD. Probing the evolutionary origins of music perception. Neurosciences and Music Ii: From Perception to Performance. New York: New York Acad Sciences; 2005. pp. 6–16.

26. Stevens C, Byron T. Universals in music processing. In: Cross I, Hallam S, Thaut M, editors. Oxford Handbook of Music Psychology. 2009. pp. 14–23.

27. Searcy WA, Nowicki S. The evolution of animal communication: reliability and deception in signaling systems. Princeton University Press; 2010.

28. Brown TJ, Handford P. Sound design for vocalizations: quality in the woods, consistency in the fields. The Condor. 2000;102: 81.

29. Lambrechts M. Organization of birdsong and constraints on performance. In: Kroodsma D, Miller E, editors. Ecology and evolution of acoustic communication in birds. Ithaca: Cornell University Press; 1996.

30. Ballentine B. Vocal performance influences female response to male bird song: an experimental test. Behav Ecol. 2004;15: 163–168.

31. Hasselquist D, Bensch S, von Schantz T. Correlation between male song repertoire, extra-pair paternity and offspring survival in the great reed warbler. Nature. 1996;381: 229–232.

32. Boogert NJ, Giraldeau LA, Lefebvre L. Song complexity correlates with learning ability in zebra finch males. Anim Behav. 2008;76: 1735–1741.

33. Farrell TM, Weaver K, An YS, MacDougall-Shackleton SA. Song bout length is indicative of spatial learning in European starlings. Behav Ecol. 2012;23: 101–111.

34. Nowicki S, Searcy WA. Song function and the evolution of female preferences - Why birds sing, why brains matter. Behavioral Neurobiology of Birdsong. New York: New York Acad Sciences; 2004. pp. 704–723.

35. Helmholtz H. On the Sensations of Tone. New York: Dover; 1885.

36. Wich SA, Nunn C. Do male “long distance calls” function in mate defense? A comparative study of long-distance calls in primates. Behav Ecol Sociobiol. 2002;52: 474–484.

37. Roederer JG. The Physics and Psychophysics of Music: An Introduction. New York: Springer; 2008.

38. Hughes R, Taylor D, Kerr R, editors. Music Lovers Encyclopedia. New York: Garden City; 1966.

39. Randel DM. The Harvard Dictionary of Music. Harvard University: Belknap Press; 2003.

40. George Dunteman. Principal components analysis. SAGE University Press; 1989.

41. Jolliffe IT. Discarding variables in a Principal Component Analysis: Artificial Data. R Stat Soc. 1972;21: 160–173.

42. Moore JM, Szekely T, Buki J, DeVoogd TJ. Motor pathway convergence predicts syllable repertoire size in oscine birds. Proc Natl Acad Sci U S A. 2011;108: 16440–16445.

43. Boogert NJ, Anderson RC, Peters S, Searcy WA, Nowicki S. Song repertoire size in male song sparrows correlates with detour reaching, but not with other cognitive measures. Anim Behav. 2011;81: 1209–1216.

44. Freedman D, Pisani R, Purves R. Statistics. 4th ed. New York: W.W. Norton & Company; 2008.

